# Assembly and Validation of Two Gap-free Reference Genomes for *Xian/indica* Rice Reveals Insights into Plant Centromere Architecture

**DOI:** 10.1101/2020.12.24.424073

**Authors:** Jia-Ming Song, Wen-Zhao Xie, Shuo Wang, Yi-Xiong Guo, Dal-Hoe Koo, Dave Kudrna, Yicheng Huang, Jia-Wu Feng, Wenhui Zhang, Yong Zhou, Andrea Zuccolo, Evan Long, Seunghee Lee, Jayson Talag, Run Zhou, Xi-Tong Zhu, Daojun Yuan, Joshua Udall, Weibo Xie, Rod A. Wing, Qifa Zhang, Jesse Poland, Jianwei Zhang, Ling-Ling Chen

**Author notes:** These authors contributed equally to this work (Jia-Ming Song, Wen-Zhao Xie, Shuo Wang). **Correspondence: Jesse Poland**, **Jianwei Zhang**, **Ling-Ling Chen**.

## Abstract

Rice (*Oryza sativa*), a major staple throughout the world and a model system for plant genomics and breeding, was the first crop genome completed almost two decades ago. However, all sequenced genomes to date contain gaps and missing sequences. Here, we report, for the first time, the assembly and analyses of two gap-free reference genome sequences of the elite *O. sativa xian/indica* rice varieties ‘Zhenshan 97 (ZS97)’ and ‘Minghui 63 (MH63)’ that are being used as a model system to study heterosis. Gap-free reference genomes also provide global insights into the structure and function of centromeres. All rice centromeric regions share conserved centromere-specific satellite motifs but with different copy numbers and structures. Importantly, we demonstrate that >1,500 genes are located in centromere regions, of which ~15.6% are actively transcribed. The generation and release of both the ZS97 and MH63 gap-free genomes lays a solid foundation for the comprehensive study of genome structure and function in plants and breed climate resilient varieties for the 21^st^ century.

## INTRODUCTION

*Oryza sativa* ‘*xian/indica*’ and ‘*geng/japonica*’ groups, previously subsp. *indica* and subsp. *japonica* respectively, are two major types of Asian cultivated rice (Wang et al., 2018). *Xian* varieties are broadly studied as they contribute over 70% of rice production worldwide and are genetically more diverse than *geng* rice. Over the past 30 years, two *xian* varieties Zhenshan 97 (ZS97) and Minghui 63 (MH63) have emerged as important model system in rice breeding and genomics being the parents of the elite hybrid Shanyou 63 (SY63), historically the most widely cultivated rice hybrid in China. Understanding the biological mechanisms behind the elite combination of ZS97 and MH63 to form the SY63 hybrid is foundational to help unravel the mystery of heterosis (Yu et al., 1997; Hua et al., 2002; Hua et al., 2003; Huang et al., 2006; Zhou et al., 2012); Further, ZS97 and MH63 represent two major varietal subgroups in *xian* rice as they show many complementary agronomic traits, and a number of important genes have been cloned based on genetic populations generated using these two varieties as parents (Sun et al., 2004; Fan et al., 2006; Xue et al., 2008). Although we previously generated two reference genome assemblies ZS97RS1 and MH63RS1 in 2016, approximately 10% of each genome remained unassembled/unplaced (Zhang et al., 2016a). Upon further analysis and editing we were able to fill the majority of gaps in each assembly and released upgraded versions of these two assemblies in 2018 (http://rice.hzau.edu.cn), yet eight (ZS97) and seven (MH63) gaps still remained.

To bridge all remaining assembly gaps across each genome we incorporated high-coverage and accurate long-reads sequence data and multiple assembly strategies to successfully generate two gap-free genome assemblies of *xian* rice ZS97 and MH63, the first gap-free plant genome assemblies publicly available to date. Importantly, we had the first opportunity to study and compare the centromeres of all chromosomes side by side across both rice varieties. More than expected, >1,500 genes were identified in rice centromere regions, ~15.6% of which were found to be actively transcribed.

## RESULTS

### Assembly and Validation of Gap-free Reference Genome Sequences for ZS97 and MH63

In this project, 56.73 Gb (~150X) and 86.85 Gb (~230X coverage) of PacBio reads (including both HiFi and CLR modes) were generated for ZS97 and MH63, respectively, using the PacBio Sequel II platform (Supplemental Figure 1, Supplemental Table 1). The PacBio HiFi and CLR reads were assembled separately with multiple *de novo* assemblers including Canu (Koren et al., 2017), FALCON (Carvalho et al., 2016), MECAT2 (Xiao et al., 2017) *etc.* (see Methods), and then the assembled contigs were merged with the two upgraded assemblies using Genome Puzzle Master (GPM) (Zhang et al., 2016b) (Supplemental Table 2-3). Finally, two gap-free reference genomes were produced, named as ZS97RS3 and MH63RS3, which contained 12 pseudomolecules with total lengths of 391.56 Mb and 395.77 Mb, respectively (Figure 1a, Table 1). Compared with the previous bacterial artificial chromosome (BAC) based RS1 genome assemblies, the new RS3 assemblies gained ~36 to 45 Mb of additional sequence by filling 223 (ZS97RS1) and 167 (MH63RS1) gaps across both genomes (Supplemental Table 4). In addition, the new assemblies corrected a few mis-orientated or mis-assembled regions caused by reliance on the Os-Nipponbare-Reference-IRGSP-1.0 sequence as a guide to produce the RS1 pseudomolecules (e.g. the 6 Mb inversion on Chr06) (Supplemental Figure 2a-c, Supplemental Table 4). These anomalies could be corrected by newly assembled contigs that were long enough to span these ambiguous regions.

**Figure 1.**
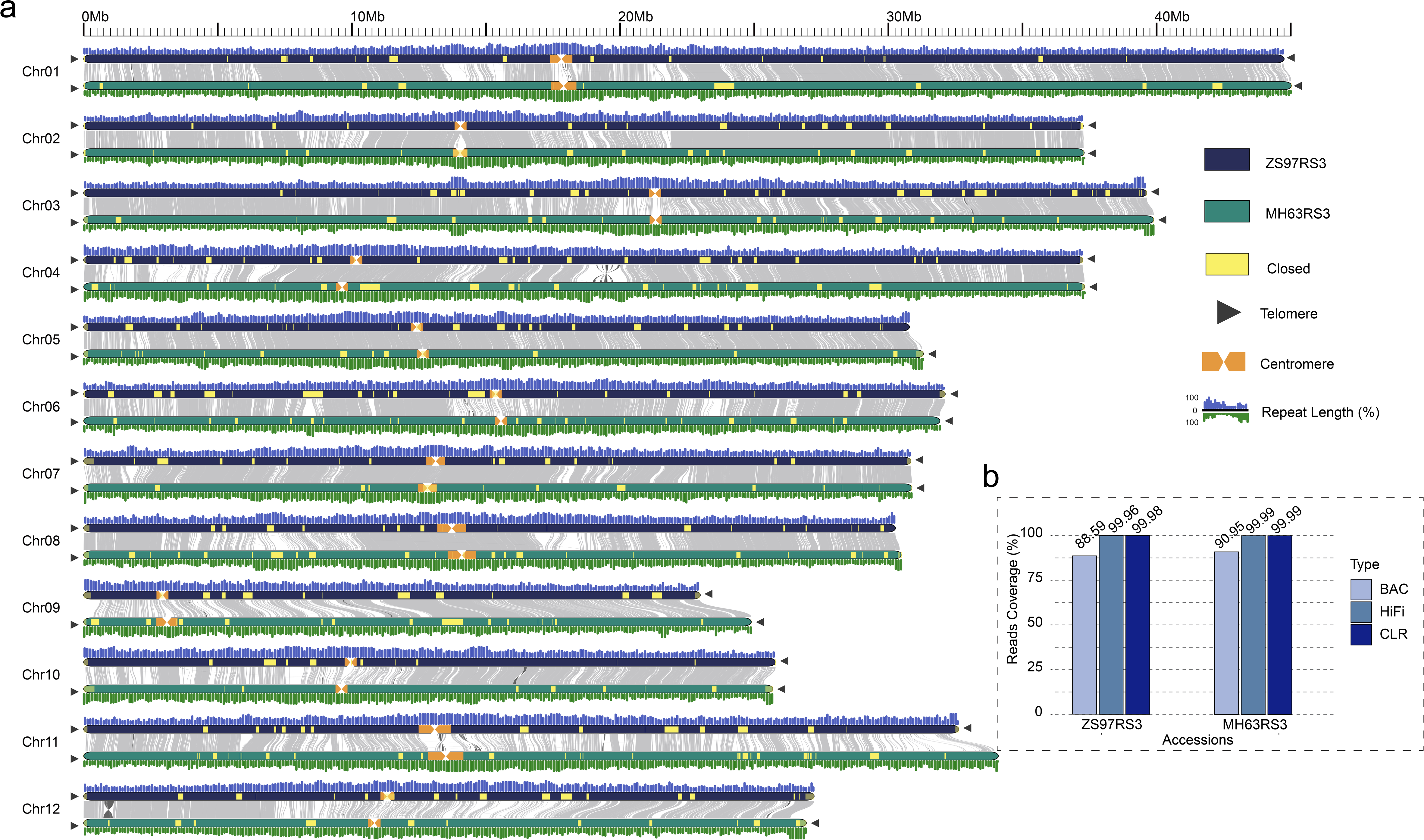
Two gap-free genomes of rice. **(A)** Collinearity analysis between ZS97RS3 and MH63RS3. The collinear regions between ZS97RS3 and MH63RS3 are shown linked by gray lines. All the RS1 gap regions closed in RS3 are showed in yellow blocks. The black triangle indicates presence of telomere sequence repeats. Repeat percentage distribution is plotted above/under each chromosome in 100-kb bins; **(B)** Histogram showed the reads coverage for different libraries in MH63RS3 and ZS97RS3, including BAC, CCS and CLR reads.

Using the 7-base telomeric repeat (CCCTAAA at 5’ end or TTTAGGG at 3’ end) as a probe, we identified 19 and 22 telomeres that resulted in 7 and 10 telomere-to-telomere (T-to-T) pseudomolecules in ZS97RS3 and MH63RS3 assemblies, respectively (Figure 1a, Supplemental Table 5-6).

The accuracy and completeness of the RS3 assemblies were validated in multiple ways. Chromosome conformation capture sequencing (Hi-C) and Bionano optical maps showed high consistency across all pseudomolecules demonstrating correct ordering and orientation (Supplemental Figure 3, Supplemental Table 2). Genome completeness was demonstrated by high mapping rates with various raw sequences, such as raw HiFi/CLR/Illumina reads, paired BAC-end sequences, and paired-end short reads from 48 RNA-seq libraries, all of which mapped at over 99% across each assembly (Supplemental Table 7-9). The evenly distributed breakpoints of aligned short and long reads indicated that all sequence connections are of high accuracy at single-base level in these final assemblies (Supplemental Figure 4). For gene content assessment, both ZS97RS3 and MH63RS3 assemblies captured 99.88% of a BUSCO 1,614 reference gene set (Supplemental Table 10). Long terminal repeat (LTR) annotation further revealed the LTR assembly index (LAI) for the ZS97RS3 and MH63RS3 assemblies were 24.01 and 22.74, respectively, which meets the standard of gold/platinum reference genomes (Ou et al., 2018, Mussurova et al., 2020) (Table 1). More than 1,500 rRNAs were identified in ZS97RS3 and MH63RS3 assemblies (Supplemental Figure 5), whereas only tens were identified in the original RS1 assemblies.

### Annotation and Comparison of Gap-free Reference Genome Sequences for ZS97 and MH63

To annotate the ZS97 and MH63 RS3 assemblies for transposable element (TE) and other repetitive sequence content, we used RepeatMasker (Zhi et al., 2006) with the latest RepBase (Bao et al., 2015) and TIGR Oryza Repeats (v3.3) (Ouyang and Buell, 2004) as libraries. As a result, we identified 465,242 TE sequences in ZS97RS3 (181.00 Mb in total length) and 468,675 TE sequences in MH63RS3 (~182.26Mb) (Supplemental Table 11-12), which accounted ~46.16% and ~45.99% of each assembly and was approximate 5% greater than that in the previous RS1 assemblies (i.e. ZS97RS1=41.28%; MH63RS3=41.58%). The repeat content increases were primarily due to the fact that over 80% of the gaps closed were in TE-rich regions (82.86% of the 45 Mb closed-gaps were TEs in ZS97RS3, and 84.17% of the 36 Mb closed-gaps were TEs in MH63RS3), and the above updated TE library.

Next we employed MAKER-P (Campbell et al., 2014) to annotate the ZS97RS3 and MH63RS3 assemblies using the identical evidence including EST, RNA-Seq, and protein used to annotate the RS1 assemblies (Supplemental Fig. 1). In order to retain consistency across different assembly versions, 51,027 and 50,341 previously annotated gene models in the ZS97RS1 and MH63RS1 assemblies, respectively, were lifted onto the RS3 annotations. Combining models annotated with MAKER-P in the newly assembled regions, the final annotations in ZS97RS3 and MH63RS3 contained 60,935 and 59,903 gene models, of which 39,258 and 39,406 were classified as non-TE gene loci (Table 1), thereby resulting in 4,648 (ZS97) and 2,082 (MH63) additional non-TE genes than previously identified in the RS1 assemblies, respectively. More than 92% of all annotated gene models were supported by homologies with known proteins or functional domains in *Oryza* and other species (Supplemental Table 13-14).

Based on our new assemblies, the annotation and comparative analyses of non-coding RNAs (transfer RNAs, ribosomal RNAs, small nucleolar RNAs, microRNAs) (Supplemental Figure 4), single nucleotide polymorphisms (SNPs) and insertions/deletions (InDels) among ZS97, MH63 and Nipponbare (Supplemental Figure 6, Supplemental Table 15), presence/absence variations (PAVs) (Supplemental Table 16), and genes in different categories (‘identical’, ‘same length’, ‘collinear’, ‘divergent’ and ‘variety-specific’ genes) (Supplemental Table 17) that were previously identified in the RS1 versions were updated.

After comparing the PAV distribution across each chromosome of both gap-free assemblies, we noticed an abundance of structural variations (SVs) near the ends of the long-arms of chromosome 11 (Figure 2a). Two large SVs, one expansion region (30.75 – 31.57 Mb) and one insertion region (31.90 – 32.76 Mb), were uniquely detected in MH63 (hereafter named as MH-E and MH-I, respectively). Raw sequencing read alignments to these two regions clearly showed that MH-E and MH-I regions could be continuously covered by MH63 reads but only partially covered by ZS97 reads (Supplemental Figure 7). Meanwhile, previous studies showed that nucleotide-binding site leucine-rich repeat (NLR) proteins were enriched in chromosome 11 (Rice Chromosomes 11 and 12 Sequencing Consortia, 2005). Hence, we performed a genome-wide homology search for NLR or NLR-like genes in both ZS97 and MH63 RS3 assemblies (Figure 2b). When putting the PAV and NLR(-like) distribution together, we could obviously determine that both MH-E and MH-I regions have more NLR(-like) content than the corresponding region in ZS97RS3 assembly (30.51 – 30.69 Mb and 30.88 – 30.94 Mb, respectively) (Supplemental Figure 7a). In the MH-E region, most of the NRL(-like) genes in ZS97 amplified 2-10 times in MH63 (Figure 2c, Supplemental Table 18), and interestingly, these genes are more likely to be expressed in root than in other tissues (Figure 2c, Supplemental Figure 7c, Supplemental Table 18). In the 857-kb MH-I region, eleven NRL(-like) genes also had higher expression levels in roots than in other tissues (Figure 2d, Supplemental Table 19). We further scanned the MH-E and MH-I homologous regions in 15 additional high-quality reference genomes (Zhou et al., 2020), and unexpectedly, none of them had both complete MH-E and MH-I at the same time (Figure 2e, Supplemental Figure 8, Supplemental Table 20). This unique genomic characteristic of MH63 could partially, at least, potentially explain its high resistance to blast disease.

**Figure 2.**
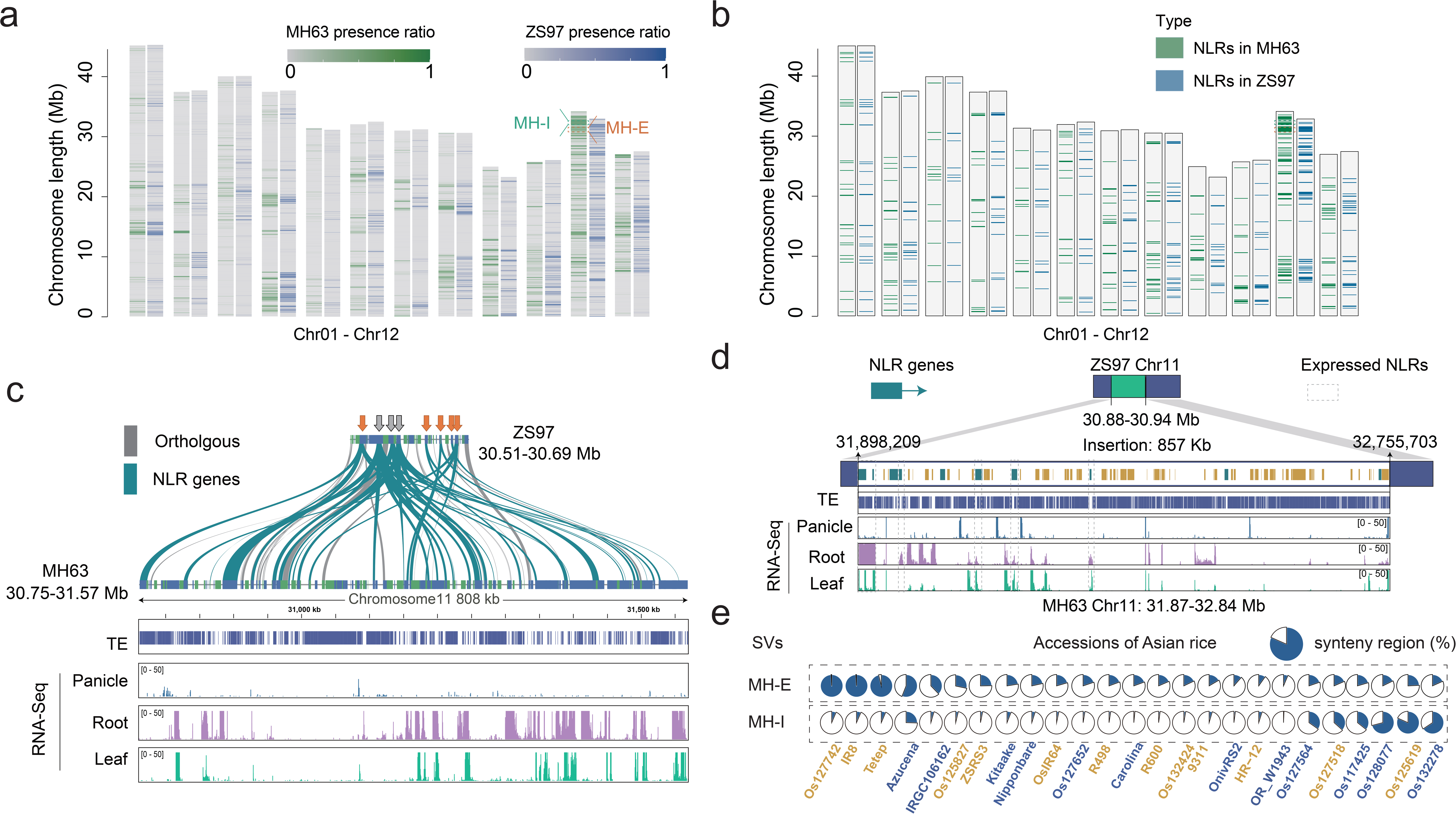
Structural variations of ZS97RS3 and MH63RS3. **(A)** Distribution of the difference regions between ZS97RS3 and MH63RS3 on the chromosome. **(B)** Distribution of the NLR genes of ZS97RS3 and MH63RS3 on the chromosome. **(C)** The expansion structural variation MH-E in MH63RS3. The structural of MH-E at the end of chromosome 11 of MH63RS3, from top to bottom are the gene collinearity of ZS97RS3 and MH63RS3, the TE distribution, the gene expression in this region. **(D)** The insertion structural variation MH-I in MH63RS3. From top to bottom are the gene collinearity of ZS97RS3 and MH63RS3, the TE distribution and the gene expression in this region. **(E)** Coverage ratio of two structural variations (MH-E and MH-I) in 25 rice varieties.

### Location and Analyses of Rice Centromeres

Centromeres are essential for maintaining the integrity of chromosomes during cell division, and ensure the fidelity of their inheritance. Unfortunately, until now, centromeres have remained largely under-explored, especially in larger genomes (Perumal et al., 2020). To functionally identify the location and sequence of centromeres in our gap-free genomes, we used the rice CENH3 antibody to immunoprecipitate chromatin from rice nuclei and then sequenced Illumina sequenced the captured DNA fragments (i.e. ChIP-Seq) (Figure 3a-b). To visually confirm the specificity of our ChIP experiments, we used fluorescent *in situ* hybridization (FISH) of ChIPed DNA on MH63 and ZS97 metaphase chromosomes, the results of which showed strong signals at the centromere for each chromosome (Figure 3b).

**Figure 3.**
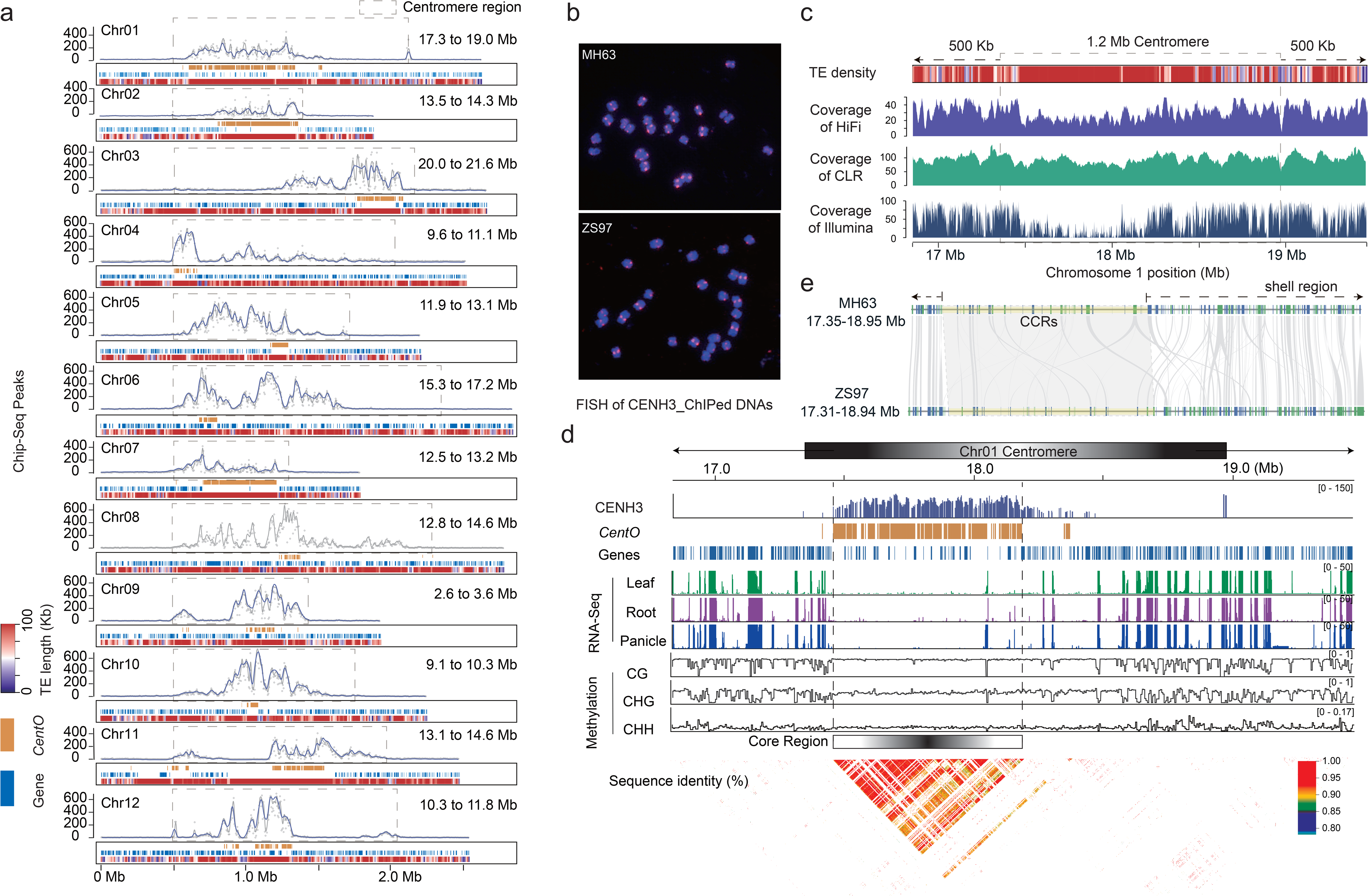
Characterization of complete rice centromeres. **(A)** The definition of MH63RS3 centromere. The layers of each chromosome graph indicate 1) the density of read mapping from CENH3 Chip-seq with sliding windows of 10-kb and 20-kb shown in grey and blue lines respectively, 2) the *CentO* satellite distribution, 3) non-TE genes distribution, and 4) TE distribution, respectively. The dotted frame represents the defined centromere area. **(B)** Fluorescence *in situ* hybridization (FISH) of mitotic metaphase chromosomes in MH63 and ZS97 using CENH3 ChIP-DNA as probe (red) with chromosomes counterstained with DAPI (blue). **(C)** Coverage of HiFi, CLR, Illumina reads and distribution of TEs in the centromere on Chr01 (extended 500 kb left and right) of MH63RS3. **(D)** The pairwise synteny visualization between ZS97RS3 and MH63RS3 in centromere area of Chr01. The synteny genes between ZS97RS3 and MH63RS3 were linked as the gray lines. The yellow blocks were core regions. **(E)** Characteristics of the centromere on Chr01 of MH63RS3. The first layer is histone CENH3 distribution, the second layer is the *CentO* distribution, the third layer is the Genes distribution, the fourth to sixth levels are gene expression, the seventh to ninth levels are methylation distribution, the tenth layer is *CentO* sequence similarity.

Using MH63RS3 as the reference, for the first time, we delimited the boundaries of each centromere and determined that the size of rice centromeres varied from 0.8 Mb to 1.8 Mb (Supplemental Figure 9, Supplemental Table 21-22). We then classified rice centromeres into core and shell regions. Core centromere regions (CCRs) were identified by sequence homology to the 155-165 bp centromere-specific (*CentO*) satellite repeats which all showed high levels of CENH3 binding (Cheng et al., 2002), while shell regions were determined by the ChIP-seq signals. The lengths of the CCRs ranged from 76 kb to 726 kb in different chromosomes with a total length of 3.47 Mb in MH63RS3 (Supplemental Figure 9, Supplemental Table 21). We manually checked the entire length of each centromere region (especially the boundary regions) of both MH63RS3 and ZS97RS3 and found that the HiFi/CLR reads were evenly mapped with no ambiguous breakpoints (Figure 3c, Supplemental Figure 10), which provides strong evidence that each of the 12 centromeres in both gap-free reference genomes were contiguous and of high quality.

Analysis results across all centromeres in both assemblies showed that CCRs contained <130 genes in each genome but a large amount of *CentO* satellite sequences (Supplemental Figure 11), while the shell regions contained >1,400 genes, of which ~16% had evidence of transcription, which included many centromere-specific retrotransposon sequences (Supplemental Tables 23-25). For example, the centromere of MH63RS3 Chr01 is 1.6 Mb, which contained a 726-kb CCR composed of 3,228 *CentO* sequences and 48 genes, while the shell regions, flanking both sides of the CCR, contained 114 *CentO* sequences and 146 genes (Figure 3d, Supplemental Table 18, Supplemental Table 23). For the genes located in CCRs of 12 chromosomes (109 in ZS97, 129 in MH63), ~10% (7 in ZS97, 13 in MH63) were found to be transcribed in the tissues and conditions tested (Supplemental Tables 24-25). Further, of the 1,446 (ZS97) and 1,764 (MH63) genes annotated in the shell regions, ~16% were found to be transcribed (231 in ZS97 and 282 in MH63). In total, 72% of gene families were shared in centromere regions of ZS97 and MH63 (Supplemental Figure 9d). Genes in the centromeres on the same chromosomes of ZS97 and MH63 were relatively conserved (mainly distributed in shell regions), an example of gene collinearity between chromosome 1 centromeres between MH63 and ZS97 was shown in Figure 3e. This conservation could be extended throughout the population structure (K=15) of cultivated Asian rice where the average ratio of conserved genes was ~87%, especially across the Chr05, Chr09 and Chr12 centromeres (Supplemental Table 26).

Gene ontology (GO) analysis showed that genes with the GO terms ‘transcription from RNA polymerase III promoter’, ‘nucleic acid binding’ and ‘nucleoplasm part’, were significantly enriched in ZS97 and MH63 centromere regions (Supplemental Figure 10b-c, Supplemental Table 27-28). Overall, these GO terms tend to have similar functions (Supplemental Figure 12). However, GO terms among different chromosomes of the same variety showed great difference, e.g., the average overlapping ratio was 37% in MH63 (Supplemental Table 29-30). We also found that the methylation levels of CG and CHG in the centromeric regions were two-fold higher than that of the whole genome (Supplemental Table 31). This phenomenon was particularly prominent in *CentO* clustered regions.

Based on the complete centromere location, we observed that the centromeric regions had slightly lower depth of mapped raw sequence reads than non-centromeric regions, which may be caused by highly repetitive elements; however, the lengths of those reads in centromeric and non-centromeric regions were broadly in line with each other (Supplemental Figure 11b). Detailed sequence analysis revealed an abundance of TEs in the centromeric region accounting for 78-80% of the functional centromere (Supplemental Table 32-33). In particular, the proportion of LTR/gypsy TEs, accounting for over 90% of the repeat content, is extremely higher than other types of TEs (Supplemental Figure 11c), which is an obvious barrier for fully assembling a centromere region.

To better understand the long-range organization and evolution of the CCRs, we generated a heat map showing pairwise sequence identity of 1-kb along the centromeres (Supplemental Figure 13a), and observed that the *CentO* sequences had the highest similarity in the middle and declined to both sides (Supplemental Figure 13a). Furthermore, the profile of *CentO* sequences (Supplemental Figure 13b) illustrated the conservation of rice centromeres on the genomic level.

To determine if the centromere architecture found in ZS97 and MH63 was conserved among other Asian rice accessions, we compared the ZS97/MH63 CCR sequences with 15 high-quality PacBio genome assemblies that represent the population structure of cultivated Asian rice (Zhou et al. 2020). The results revealed that the lengths of *CentO* satellite repeats in the CCRs of the same chromosomes varied significantly between populations, and varieties within the same populations (Supplemental Table 34-35).

## DISCUSSION

In this study, we assembled and validated the first two gap-free reference genome sequences of rice available to the research community. At present, this work could only have been achieved with a combination of multiple and deep-coverage sequence datasets, cutting-edge technologies and assemblers, verses reliance on a single sequence dataset and assembler. For example, none of the *de novo* assemblers could ideally produce all complete pseudomolecules for the 12 rice chromosomes, but a set of fragmented contigs (Supplemental Table 3). Even with the same dataset, assembly results varied when using different assemblers and parameters. As we observed in our project, the data obtained by different sequencing approaches have different coverages: i.e. both the PacBio HiFi and CLR reads covered >99.9% of the ZS97RS3 and MH63RS3 gap-free genomes, while BAC-based reads of RS1 assemblies only covered 88.59% and 90.95%, respectively (Figure 1b). Hence, the strategy applied here was to fully leverage the complementarity of datasets, assemblers and technologies. In our final assemblies, we manually selected and merged proper sequence contigs to cover their corresponding regions across each chromosome (Supplemental Figure 6). The last closed gaps in our assemblies were all in centromere regions, which confirms that the great challenge for completely assembling plant genomes is was from the nature of their complicated architecture and highly repetitive sequences. Long-read sequencing data of high accuracy, however, can span the repeats allowed assemblers to distinguish minor sequence differences in repeat regions during the assembling process.

We also used independent methods like Hi-C and Bionano maps to validate our assemblies, as well as FISH and ChIP-Seq assays to discover and characterize the location and primary structure of centromeres.

In conclusion, the generation and validation of two gap-free assemblies of ZS97 and MH63, presented here, provides a clear picture of the primary sequence architecture of the *xian/indica* rice genomes that feed the world. Such data will serve as a fundamental and comprehensive model resource in the study of hybrid vigor, and other basic and applied research, and leads the path forward to a new standard for reference genomes in plant biology.

## METHODSs

### Plant Materials and Sequencing

Fresh young leaf tissue was collected from *O. sativa* ZS97 and MH63 plants. We constructed SMRTbell libraries as described in previous study (Pendletonet al., 2015). The genomes of MH63 and ZS97 were sequenced using the PacBio Sequel II platform (Pacific Biosciences), to produce 8.34 Gb HiFi reads (~23x coverage) and 48.39Gb CLR reads (~131x coverage) for ZS97, and 37.88 Gb HiFi reads (~103x coverage) and 48.97 Gb CLR reads (~132x coverage) for MH63 genomes.

Truseq Nano DNA HT Sample preparation Kit following manufacturer’s standard protocol (Illumina) was used to generate the libraries for Illumina paired-end genome sequencing. These libraries were sequenced to generate 150 bp paired-end reads by Illumina HiSeq X Ten platform with 350 bp insert size, and produce 25 Gb reads (~69x coverage) for ZS97, and 28 Gb reads (~76x coverage) for MH63 genomes.

Plant tissues used for optical mapping were extracted using the Bionano plant tissue extraction protocol (Staňková et al., 2016). Extracted DNA was embedded in BioRad LE agarose for subsequent washes of T.E., proteinase K (0.8mg/ml), and RNAse A (20μL/mL) treatments in lysis buffer. The agarose plugs were then melted using agarase (0.1 U/μL, New England Biolabs) and dialyzed on millipore membranes (0.1μm) with T.E. to equilibrate ion concentrations. The DNA was then nicked with the nickase restriction enzyme BssSI (2U/μL). Labeled nucleotides were incorporated at breakpoints and the DNA was counterstained. Each sample was loaded onto 2 nanochannel flow cells of a Bionano Irys machine for DNA imaging.

### Genome Assembly and Assessment

Seven tools based on different algorithms were used to assemble the genomes of ZS97 and MH63. (1) Canu v1.8 (Koren et al., 2017) was used to assemble the genomes with default parameters; (2) FALCON toolkit v0.30 (Carvalho et al., 2016) was applied for assembly with the parameters: pa_DBsplit_option = -s200 -x500, ovlp_DBsplit_option = -s200 -x500, pa_REPmask_code = 0,300;0,300;0,300, genome_size = 400000000, seed_coverage = 30, length_cutoff = -1, pa_HPCdaligner_option = -v -B128 -M24, pa_daligner_option = -k18 -w8 -h480 -e.80 -l5000 -s100, falcon_sense_option = --output-multi --min-idt 0.70 --min- cov 3 --max-n-read 400, falcon_sense_greedy = False, ovlp_HPCdaligner_option = -v -M24 -l500, ovlp_daligner_option = -h60 -e0.96 -s1000, overlap_filtering_setting = --max-diff 100 --max-cov 100- -min-cov 2, length_cutoff_pr = 1000; (3) MECAT2 (Xiao et al., 2017) was utilized to assemble with the parameters: GENOME_SIZE = 400000000, MIN_READ_LENGTH = 2000, CNS_OVLP_OPTIONS = “”, CNS_OPTIONS = “-r 0.6 -a 1000 -c 4 -l 2000”, CNS_OUTPUT_COVERAGE = 30, TRIM_OVLP_OPTIONS = “-B”, ASM_OVLP_OPTIONS = “-n 100 -z 10 -b 2000 -e 0.5 -j 1 -u 0 -a 400”, FSA_OL_FILTER_OPTIONS = “--max_overhang = -1 --min_identity = - 1”, FSA_ASSEMBLE_OPTIONS = “”, GRID_NODE = 0, CLEANUP = 0, USE_GRID = false; (4) Flye 2.6-release (Kolmogorov et al., 2019) was set with “--genome-size 400m”; (5) Wtdbg2 2.5 (Ruan and Li., 2020) was used to assemble with parameters “-x sq, -g 400m”, and then Minimap2 (Li, 2018) was employed to map the PacBio CLR data to the assembly results, and wtpoa was utilized polish and correct the wtdbg2 assembly results; (6) NextDenovo v2.1-beta.0 (https://github.com/Nextomics/NextDenovo) was applied for assembly with parameters “task = all, rewrite = yes, deltmp = yes, rerun = 3, input_type = raw, read_cutoff = 1k, seed_cutoff = 44382, blocksize = 2g, pa_correction = 20, seed_cutfiles = 20, sort_options = -m 20g -t 10 -k 40, minimap2_options_raw = -x ava-ont -t 8, correction_options = -p 10, random_round = 20, minimap2_options_cns = -x ava-pb -t 8 -k17- w17, nextgraph_options = -a 1”; (7) Miniasm-0.3-r179 (Li, 2016) with default parameters.

Based on the results of these seven software tools, Genome Puzzle Master (GPM) (Zhang et al., 2016b) was then used to integrate and optimize the assembled contigs, and visualize complete chromosomes. Based on the HiFi and CLR sequencing data, we used the GenomicConsensus package of SMRTLink/7.0.1.66975 (https://www.pacb.com/support/) to polish the assembled genomes twice with the Arrow algorithm, using the parameter: --algorithm=arrow. Pilon (Walker et al., 2014) was used for polishing the genomes based on Illumina data with the parameters: --fix snps, indels. This process was repeated twice. Molecules were then assembled using Bionano IrysSolve pipeline (https://bionanogenomics.com/support-page/) to create optical maps. Images were interpreted quantitatively using Bionano AutoDetect 2.1.4.9159 and data was visualized using IrysView v2.5.1. These assemblies were used with draft genome assemblies to validate and scaffold the sequences. Bionano optical map data was aligned to the merged contigs using RefAlignerAssembler in the IrysView software package to perform the verification.

ZS97RS3 and MH63RS3 genome completeness was assessed using BUSCO v4.0.6, which contained 1614 genes in the ‘embryophyta_odb10’ dataset (Simão et al., 2015), with default parameters. In addition, we mapped the PacBio HiFi reads and PacBio CLR reads with Minimap2 (Li, 2018), Illimina reads with BWA-0.7.17 (Jo et al., 2015), BES/BAC reads with BLASTN v2.7.1 (Altschul et al., 1990), Hi-C reads with HiC-Pro v2.11.1 (Servant et al., 2015), RNA-Seq reads with Hisat2 v2.1.0 (Kim et al., 2015) to both genome assemblies.

### Gene and Repeat Annotations

MAKER-P (Campbell et al., 2014) version 3 was used to annotate the ZS97RS3 and MH63RS3 genomes. All evidence was the same as that used for RS1 genome annotations. To ensure the consistency with the RS1 versions, genes that mapped in the entirety to the RS3 genomes were retained. New genes in gap regions were obtained from MAKER-P results (Campbell et al., 2014). Genes encoding transposable elements were identified and transitively annotated by searching against the MIPS-REdat Poaceae version 9.3p (Nussbaumer et al., 2013) database using TBLASTN (Altschul et al., 1990) with an E-value of 1e-10. tRNAs were identified with tRNAscan-SE (Lowe and Eddy, 1997) using default parameters; rRNA genes were identified by searching each assembly against the rRNA sequences of Nipponbare using BLASTN v2.7.1 (Altschul et al., 1990); miRNAs and snRNAs were predicted using INFERNAL of Rfam (Griffiths-Jones et al., 2005) (v14.1). Repeats in the genome were annotated using RepeatMasker (Zhi et al., 2006) with RepBase (Bao et al., 2015), and TIGR Oryza Repeats (v3.3) with RMBlast search engine. For the overlapping repeats in different classes, LTR retrotransposons were kept first, next TIR, and then SINE and LINE, finally helitrons. This priority order was based on stronger structural signatures. Besides, the known nested insertions models (LTR into helitron, helitron into LTR, TIR into LTR, LTR into TIR) were retained. The identified repetitive elements were further characterized and classified using PGSB repeat classification schema. LTR_FINDER (Xu and Wang 2007) was used to identify complete LTR-RTs with target site duplications (TSDs), primer binding sites (PBS) and polypurine tract (PPT).

### Chromatin Immunoprecipitation (ChIP) and ChIP-seq

Procedures for chromatin immunoprecipitation (ChIP) were adopted from Nagaki et al. (2003) and Walkowiak et al. (2020) using nuclei from 4-week-old seedlings. Chromatin with the nuclei was digested with micrococcal nuclease (Sigma-Aldrich, St. Louis, MO) to liberate nucleosomes. For ChIP, we used anti-centromeric histone 3 antibody (N-term) which reacts with 18.5 kDa CenH3 protein from Oryza sativa purchased from Antibodies-online Inc. (Limerick, PA; cat# ABIN1106669). The digested mixture was then incubated overnight with 3 μg of rice CENH3 antibody at 4°C. The target antibodies were then captured from the mixture using Dynabeads Protein G (Invitrogen, Carlsbad, CA). ChIP-seq libraries were then constructed using a TruSeq ChIP Library Preparation Kit (Illumina, San Diego, CA) following the manufacturer’s instructions and the libraries were sequenced (2×150bp) on an Illumina HiSeqX10.

### Fluorescence *in situ* Hybridization (FISH)

#### Slide Preparation

Mitotic chromosomes were prepared as described by Koo and Jiang (2009) with minor modifications. Root tips were collected from plants and treated in a nitrous oxide gas chamber for 1.5 h. The root tips were fixed overnight in ethanol:glacial acetic acid (3:1) and then squashed in a drop of 45% acetic acid.

#### Probe Labeling and Detection

The ChIPed DNAs were labeled with digoxigenin-16-dUTP using a nick translation reaction. The clone, maize 45S rDNA (Koo and Jiang 2009) was labeled with biotin-11-dUTP (Roche, Indianapolis, IN). Biotin- and digoxigenin-labeled probes were detected with Alexa Fluor 488 streptavidin antibody (Invitrogen) and rhodamine-conjugated anti-digoxigenin antibody (Roche), respectively. Chromosomes were counterstained with 4’,6-diamidino-2-phenylindole (DAPI) in Vectashield antifade solution (Vector Laboratories, Burlingame, CA). Images were captured with a Zeiss Axioplan 2 microscope (Carl Zeiss Microscopy LLC, Thornwood, NY) using a cooled CCD camera CoolSNAP HQ2 (Photometrics, Tucson, AZ) and AxioVision 4.8 software. The final contrast of the images was processed using Adobe Photoshop CS5 software.

### The Completeness of Centromeres on MH63RS3 and ZS97RS3 Chromosomes

Based on the final RS3 genome assemblies, we use BLAST (Altschul et al., 1990) to align the *CentO* satellite repeats in rice to each reference genome with an E-value of 1e-5, and then use BEDtools (Quinlan, 2014) to merge the results with the parameter ‘-d 50000’. If a region contained more than 10 consecutive *CentO* repeats with lengths longer than 10 kb, it was classified as core centromere region (CCR).

For the identification of the whole centromere region, we use BWA-0.7.17 (Jo and Koh., 2015) to align the CENH3 ChIP-Seq reads to MH63RS3 and ZS97RS3 genomes, and use SAMtools (Li et al., 2009) to filter the results with mapQ value above 30; then we use MACS2 (Zhang et al., 2008) to call the peaks of CENH3. Finally, we manually defined all the centromeric region of MH63RS3 and ZS97RS3 genomes in consideration of the distribution of CENH3 histone, *CentO*, repeats and genes.

To compare of *CentO* sequence similarity, we first used BEDtools (Quinlan, 2014) to obtain sequences of centromere core regions, and divide them into 1 kb continuous sequences; then we used Minimap2 (Li, 2018) to align the sequences with the parameters: -f 0.00001 -t 8 -X --eqx -ax ava -pb; and, finally, used a custom python script to filter the results file, and used R to generate a heat map showing pairwise sequence identity (Logsdon et al., 2020).

### Telomere Sequence Identification

The telomere sequence 5’-CCCTAAA-3’ and the reverse complement of the seven bases were searched directly. In addition, we used BLAT (Kent, 2002) to search telomere-associated tandem repeats sequence (TAS) from TIGR *Oryza* Repeat database (Ouyang and Buell, 2004) in whole genome.

### Identification of PAVs between ZS97RS3 and MH63RS3

ZS97RS3 assembly was aligned to MH63RS3 assembly using Mummer (4.0.0beta2) (Marçais et al., 2018) with parameters settings ‘-c 90 -l 40’. Then we used “delta-filter −1” parameter with the one-to-one alignment block option to filter the alignment results. Further “show-diff” was used to select for unaligned regions as the PAVs.

### Prediction of NLR Genes

We first predicted domains of genes with InterProScan (Jones et al., 2014), which can analyze peptide sequences against InterPro member databases, including ProDom, PROSITE, PRINTS, Pfam, PANTHER, SMART and Coils. Pfam and Coils were used to prediction NLRs which were required to contain at least one NB, TIR, or CC_R_ (RPW8) using the following reference sequences: NB (Pfam accession PF00931), TIR (PF01582), RPW8 (PF05659), LRR (PF00560, PF07725, PF13306, PF13855) domains, or CC motifs (Van de Weyer et al., 2019).

### Identification of Collinear Orthologues

MCscan (python version) (Tang et al., 2008) was used to identify collinear orthologues between chromosome 11 of ZS97RS3 and MH63RS3 genomes with default parameters.

## DATA AVAILABILITY

All the raw sequencing data generated for this project are achieved at NCBI under accession numbers SRR13280200, SRR13280199 and SRR13288213 for ZS97, SRX6957825, SRX6908794, SRX6716809 and SRR13285939 for MH63. The genome assemblies are available at NCBI (CP056052-CP056064 for ZS97RS3, CP054676-CP054688 for MH63RS3) and annotations are visualized with Gbrowse at http://rice.hzau.edu.cn. All the materials in this study are available upon request.

## FUNDING

This research was supported by the National Key Research and Development Program of China (2016YFD0100904 and 2016YFD0100802), the National Natural Science Foundation of China (31871269), Hubei Provincial Natural Science Foundation of China (2019CFA014), the Fundamental Research Funds for the Central Universities (2662020SKPY010 to J.Z.).

## AUTHOR CONTRIBUTIONS

L.-L.C., J.Z., R.W. and Q.Z. designed studies and contributed to the original concept of the project. J.P. and D.-H.K. performed the ChIP-seq and FISH experiments. D.K., E.L., S.L., J.T., D.Y., J.U. and R.W. performed the genome and Bionano sequencing. J.-M.S., W.-Z.X., S.W., Y.-X.G., Y.H. J.-W.F., W.Z., R.Z. and X.T.Z. performed genome assembling and annotation, comparative genomics analysis and other data analysis. J.-M.S., W.-Z.X., S.W., J.P., D.-H.K., L.-L.C. and J.Z. wrote the paper. W.X., R.W. and Q.Z. contributed to revisions.

## ACKNOWLEDGEMENTS

We sincerely thank 1) Pacific Biosciences of California, Inc. for sequencing of MH63, 2) Wuhan Frasergen Bioinformatics Co., Ltd. for sequencing of ZS97 and 3) Dr. Jiming Jiang at MSU for his critical comments and constructive suggestions on our centromere analyses.

## ONLINE CONTENT

Any methods, additional references, research reporting summaries, source data, statements of code and data availability and associated accession codes are available online.

## Notes

### Competing Interest Statement

The authors have declared no competing interest.

## REFERENCES

Altschul, S.F., Gish, W., Miller, W., Myers, E.W., and Lipman, D.J. (1990). Basic local alignment search tool. J. Mol. Biol. 215: 403–410.

Bao, W., Kojima, K.K., and Kohany, O. (2015). Repbase update, a database of repetitive elements in eukaryotic genomes. Mob. DNA 6: 11.

Campbell, M.S., Law, M., Holt, C., Stein, J.C., Moghe, G.D., Hufnagel, D.E., Lei, J., Achawanantakun, R., Jiao, D., Lawrence, C.J. et al. (2014). MAKER-P: a tool kit for the rapid creation, management, and quality control of plant genome annotations. Plant Physiol. 164: 513–524.

Carvalho, A.B., Dupim, E.G., and Goldstein, G. (2016). Improved assembly of noisy long reads by k-mer validation. Genome Res. 26: 1710–1720.

Cheng, Z., Dong, F., Langdon, T., Ouyang, S., Buell, C.R., Gu, M., Blattner, F.R., and Jiang, J. (2002). Functional rice centromeres are marked by a satellite repeat and a centromere-specific retrotransposon. Plant Cell 14: 1691–1704.

Simão, F.A., Waterhouse R.M., Ioannidis P., Kriventseva E.V., and Zdobnov, E.M. (2015). BUSCO: assessing genome assembly and annotation completeness with single-copy orthologs. Bioinformatics 31: 3210–3212.

Fan, C., Xing, Y., Mao, H., Lu, T., Han, B., Xu, C., Li, X., and Zhang, Q. (2006). GS3, a major QTL for grain length and weight and minor QTL for grain width and thickness in rice, encodes a putative transmembrane protein. Theor. Appl. Genet. 112: 1164–1171.

Fu, L., Niu, B., Zhu, Z., Wu, S., and Li, W. (2012). CD-HIT: accelerated for clustering the next generation sequencing data. Bioinformatics 28: 3150–3152.

Griffiths-Jones, S., Moxon, S., Marshall, M., Khanna, A., Eddy, S.R., and Bateman, A. (2005). Rfam: annotating non-coding RNAs in complete genomes. Nucleic Acids Res. 33: D121–D124.

Hua, J., Xing, Y., Wu, W., Xu, C., Sun, X., Yu, S., and Zhang, Q. (2003). Single-locus heterotic effects and dominance by dominance interactions can adequately explain the genetic basis of heterosis in an elite rice hybrid. Proc. Natl. Acad. Sci. USA 100: 2574–2579.

Hua, J.P., Xing, Y.Z., Xu, C.G., Sun, X.L., Yu, S.B., and Zhang, Q. (2002). Genetic dissection of an elite rice hybrid revealed that heterozygotes are not always advantageous for performance. Genetics 162: 885–1895.

Huang, Y., Zhang, L., Zhang, J., Yuan, D., Xu, C., Li, X., Zhou, D., Wang, S., and Zhang, Q. (2006). Heterosis and polymorphisms of gene expression in an elite rice hybrid as revealed by a microarray analysis of 9198 unique ESTs. Plant Mol. Biol. 62: 579–591.

Jo, H., and Koh, G. (2015). Faster single-end alignment generation utilizing multi-thread for BWA. Biomed Mater Eng. Suppl 1: S1791–1796.

Jones, P., Binns, D., Chang, H.Y., Fraser, M., Li, W., McAnulla, C., McWilliam, H., Maslen, J., Mitchell, A., Nuka, G., et al. (2014). InterProScan 5: genome-scale protein function classification. Bioinformatics 30: 1236–1240.

Kent, W.J. (2002). BLAT--the BLAST-like alignment tool. Genome Res. 12: 656–664.

Kim, D., Langmead, B., and Salzberg, S.L. (2015). HISAT: a fast spliced aligner with low memory requirements. Nat. Methods 12:357–360.

Kolmogorov, M., Yuan, J., Lin, Y., and Pevzner, P.A. (2019). Assembly of long, error-prone reads using repeat graphs. Nat. Biotechnol. 37: 540–546.

Koo, D.H., and Jiang, J.M. (2009). Super-stretched pachytene chromosomes for fluorescence in situ hybridization mapping and immunodetection of cytosine methylation. Plant J. 59: 509–516.

Koren, S., Walenz, B.P., Berlin, K., Miller, J.R., Bergman, N.H., and Phillippy, A.M. (2017). Canu: scalable and accurate long-read assembly via adaptive k-mer weighting and repeat separation. Genome Res. 27: 722–736.

Li, H. (2016). Minimap and miniasm: fast mapping and de novo assembly for noisy long sequences. Bioinformatics 32: 2103–2110.

Li, H. (2018). Minimap2: pairwise alignment for nucleotide sequences. Bioinformatics 34: 3094–3100.

Li, H., and Durbin, R. (2009). Fast and accurate short read alignment with Burrows-Wheeler transform. Bioinformatics 25: 1754–1760.

Li, H., Handsaker, B., Wysoker, A., Fennell, T., Ruan, J., Homer, N., Marth, G., Abecasis, G., Durbin, R., and 1000 Genome Project Data Processing Supgroup. (2009). The Sequence Alignment/Map format and SAMtools. Bioinformatics 25: 2078–2079.

Logsdon, G.A., Vollger, M.R., Hsieh, P.H., Mao, Y., Liskovykh, M.A., Koren, S., Nurk, S., Mercuri, L., Dishuck, P.C., Rhie, A., et al. (2020). The structure, function, and evolution of a complete human chromosome 8. bioRxiv 2020.09.08.285395

Lowe, T.M., and Eddy, S.R. (1997). tRNAscan-SE: a program for improved detection of transfer RNA genes in genomic sequence. Nucleic Acids Res. 25: 955–964.

Marçais, G., Delcher, A.L., Phillippy, A.M., Coston, R., Salzberg, S.L., and Zimin, A. (2018). MUMmer4: A fast and versatile genome alignment system. PLoS Comput. Biol. 14: e1005944.

Miga, K.H., Koren, S., Rhie, A., Vollger, M.R., Gershman, A., Bzikadze, A., Brooks, S., Howe, E., Porubsky, D., Logsdon, G.A., et al. (2020). Telomere-to-telomere assembly of a complete human X chromosome. Nature 585: 79–84.

Mussurova, S., Al-Bader, N., Zuccolo, A., and Wing, R.A. (2020). Potential of platinum standard reference genomes to exploit natural variation in the wild relatives of rice. Front Plant Sci. 11: 579980.

Nagaki, K., Talbert, P.B., Zhong, C.X., Dawe, R.K., Henikoff, S., and Jiang, J. (2003). Chromatin immunoprecipitation reveals that the 180-bp satellite repeat is the key functional DNA element of Arabidopsis thaliana centromeres. Genetics 163: 1221–1225.

Nussbaumer, T., Martis, M.M., Roessner, S.K., Pfeifer, M., Bader, K.C., Sharma, S., Gundlach, H., and Spannagl, M. (2013). MIPS PlantsDB: a database framework for comparative plant genome research. Nucleic Acids Res. 41: D1144–1151.

Ou, S., Chen, J., and Jiang, N. (2018). Assessing genome assembly quality using the LTR Assembly Index (LAI). Nucleic Acids Res. 46: e126.

Ouyang, S., and Buell, C.R. (2004). The TIGR Plant Repeat Databases: a collective resource for the identification of repetitive sequences in plants. Nucleic Acids Res. 32: D360–363.

Pierce, N.T., Irber, L., Reiter, T., Brooks, P., and Brown, C.T. (2019). Large-scale sequence comparisons with *sourmash*. F1000Res 8: 1006.

Pendleton, M., Sebra, R., Pang, A.W., Ummat, A., Franzen, O., Rausch, T., Stütz, A.M., Stedman, W., Anantharaman, T., Hastie, A., et al. (2015). Assembly and diploid architecture of an individual human genome via single-molecule technologies. Nat. Methods 12: 780–786.

Perumal, S., Koh, C.S., Jin, L., Buchwaldt, M., Higgins, E.E., Zheng, C., Sankoff, D., Robinson, S.J., Kagale, S., Navabi, Z.K., et al. (2020). A high-contiguity *Brassica nigra* genome localizes active centromeres and defines the ancestral Brassica genome. Nat Plants 6: 929–941.

Quinlan, A.R. (2014). BEDTools: the swiss-army tool for genome feature analysis. Curr. Protoc. Bioinformatics 47: 11.12.134.

Rice Chromosomes 11 and 12 Sequencing Consortia. (2005). The sequence of rice chromosomes 11 and 12, rich in disease resistance genes and recent gene duplications. BMC Biol. 3: 20.

Ruan, J., and Li, H. (2020). Fast and accurate long-read assembly with wtdbg2. Nat. Methods 17: 155–158.

Servant, N., Varoquaux, N., Lajoie, B.R., Viara, E., Chen, C.J., Vert, J.P., Heard, E., Dekker, J., and Barillot, E. (2015). HiC-Pro: an optimized and flexible pipeline for Hi-C data processing. Genome Biol. 16:259.

Simão, F.A., Waterhouse, R.M., Ioannidis, P., Kriventseva, E.V., Zdobnov, E.M. (2015). BUSCO: assessing genome assembly and annotation completeness with single-copy orthologs. Bioinformatics 31: 3210–3212.

Staňková, H., Hastie, A.R., Chan, S., Vrána, J., Tulpová, Z., Kubaláková, M., Visendi, P., Hayashi, S., Luo, M., Batley, J., et al. (2016). BioNano genome mapping of individual chromosomes supports physical mapping and sequence assembly in complex plant genomes. Plant Biotechnol J. 14: 1523–1531.

Sun, X., Cao, Y., Yang, Z., Xu, C., Li, X., Wang, S., and Zhang, Q. (2004) Xa26, a gene conferring resistance to *Xanthomonas oryzae* pv. *oryzae* in rice, encodes an LRR receptor kinase-like protein. Plant J. 37: 517–527.

Tang, H., Bowers, J.E., Wang, X., Ming, R., Alam, M., and Paterson, A.H. (2008). Synteny and collinearity in plant genomes. Science 320: 486–488.

Van de Weyer, A.-L., Monteiro, F., Furzer, O.J., Nishimura, M.T., Cevik, V., Witek, K., Jones, J.D.G., Dangl, J.L., Weigel, D., and Bemm, F. (2019). A species-wide inventory of NLR genes and alleles in Arabidopsis thaliana. Cell 178:1260–1272.

Walkowiak, S., Gao, L., Monat, C., Haberer, G., Kassa, M.T., Brinton, J., Ramirez-Gonzalez, R.H., Kolodziej, M.C., Delorean, E., Thambugala, D., et al. (2020). Multiple wheat genomes reveal global variation in modern breeding. Nature 588: 277–283.

Walker, B.J., Abeel, T., Shea, T., Priest, M., Abouelliel, A., Sakthikumar, S., Cuomo, C.A., Zeng, Q., Wortman, J., Young, S.K., and Earl, A.M. (2014). Pilon: an integrated tool for comprehensive microbial variant detection and genome assembly improvement. PLoS One 9:e112963.

Wang, W., Mauleon, R., Hu, Z., Chebotarov, D., Tai, S., Wu, Z., Li, M., Zheng, T., Fuentes, R.R., Zhang, F., et al. (2018). Genomic variation in 3,010 diverse accessions of Asian cultivated rice. Nature 557: 43–49.

Xiao, C.L., Chen, Y., Xie, S.Q., Chen, K.N., Wang, Y., Han, Y., Luo, F., and Xie, Z. (2017). MECAT: fast mapping, error correction, and de novo assembly for single-molecule sequencing reads. Nat. Methods 14: 1072–1074.

Xue, W., Xing, Y., Weng, X., Zhao, Y., Tang, W., Wang, L., Zhou, H., Yu, S., Xu, C., Li, X., and Zhang, Q. (2008) Natural variation in Ghd7 is an important regulator of heading date and yield potential in rice. Nat. Genet. 40: 761–767.

Xu, Z., and Wang, H. (2007). LTR_FINDER: an efficient tool for the prediction of full-length LTR retrotransposons. Nucleic Acids Res. 35: W265–268.

Yu, S.B., Li, J.X., Xu, C.G., Tan, Y.F., Gao, Y.J., Li, X.H., Zhang, Q., and Saghai Maroof, M.A. (1997). Importance of epistasis as the genetic basis of heterosis in an elite rice hybrid. Proc. Natl. Acad. Sci. USA 94: 9226–9231.

Zhang, J., Chen, L.L., Xing, F., Kudrna, D.A., Yao, W., Copetti, D., Mu, T., Li, W., Song, J.M., Xie, W., et al. (2016a). Extensive sequence divergence between the reference genomes of two elite indica rice varieties Zhenshan 97 and Minghui 63. Proc. Natl. Acad. Sci. USA 113: E5163–5171.

Zhang, J., Kudrna, D., Mu, T., Li, W., Copetti, D., Yu, Y., Goicoechea, J.L., Lei, Y., and Wing, R.A. (2016b). Genome puzzle master (GPM): an integrated pipeline for building and editing pseudomolecules from fragmented sequences. Bioinformatics 32: 3058–3064.

Zhang, Y., Liu, T., Meyer, C.A., Eeckhoute, J., Johnson, D.S., Bernstein, B.E., Nusbaum, C., Myers, R.M., Brown, M., Li, W., and Liu, X.S. (2008). Model-based analysis of ChIP-Seq (MACS). Genome Biol. 9: R137.

Zhi, D., Raphael, B.J., Price, A.L., Tang, H., and Pevzner, P.A. (2006). Identifying repeat domains in large genomes. Genome Biol. 7: R7.

Zhou, G., Chen, Y., Yao, W., Zhang, C., Xie, W., Hua, J., Xing, Y., Xiao, J., and Zhang, Q. (2012). Genetic composition of yield heterosis in an elite rice hybrid. Proc. Natl. Acad. Sci. USA 109: 15847–15852.

Zhou, Y., Chebotarov, D., Kudrna, D., Llaca, V., Lee, S., Rajasekar, S., Mohammed, N., Al-Bader, N., Sobel-Sorenson, C., Parakkal, P., et al. (2020). A platinum standard pan-genome resource that represents the population structure of Asian rice. Sci. Data 7: 113.

